# Wiring together large single-cell RNA-seq sample collections

**DOI:** 10.1101/460246

**Authors:** Nikolas Barkas, Viktor Petukhov, Daria Nikolaeva, Yaroslav Lozinsky, Samuel Demharter, Konstantin Khodosevich, Peter V. Kharchenko

**Affiliations:** Department of Biomedical Informatics, Harvard Medical School, Boston, MA 02115 USA; Biotech Research and Innovation Centre (BRIC), Faculty of Health, University of Copenhagen, Copenhagen N, DK-2200 Denmark; Harvard Stem Cell Institute, Cambridge, MA 02315 USA

## Abstract

Single-cell RNA-seq methods are being increasingly applied in complex study designs, which involve measurements of many samples, commonly spanning multiple individuals, conditions, or tissue compartments. Joint analysis of such extensive, and often heterogeneous, sample collections requires a way of identifying and tracking recurrent cell subpopulations across the entire collection. Here we describe a flexible approach, called Conos (Clustering On Network Of Samples), that relies on multiple plausible inter-sample mappings to construct a global graph connecting all measured cells. The graph can then be used to propagate information between samples and to identify cell communities that show consistent grouping across broad subsets of the collected samples. Conos results enable investigators to balance between resolution and breadth of the detected subpopulations. In this way, it is possible to focus on the fine-grained clusters appearing within more similar subsets of samples, or analyze coarser clusters spanning broader sets of samples in the collection. Such multi-resolution joint clustering provides an important basis for downstream analysis and interpretation of sizeable multi-sample single-cell studies and atlas-scale collections.

---

Until recently, joint consideration of multiple single-cell RNA-seq (scRNA-seq) datasets has focused on analysis of technical or biological replicates that were being generated to accrue more cells and ensure consistency^1-3^. In such settings, the joint analysis can be formulated as a batch correction problem, relying on an assumption that the underlying sample composition is nearly identical, and treating inter-sample variation as a technical artifact that needs to be controlled for^4,5^. Recent works by Buttler *et al*^6^ and Haghverdi *et al*^7^ have introduced more general methods for aligning multiple scRNA-seq samples, allowing for greater differences between the samples. While these new methods still aimed to bring multiple datasets to a common set of expression coordinates where matching subpopulations from different samples would be intermixed, they were able to handle notably more diverse samples. This, for instance, enabled alignment of datasets that did not share all of the cell subpopulations, datasets generated using different platforms, or even datasets originating from different species. The two methods, nevertheless, were designed to align relatively small sample panels with modest compositional variation.

The progress of scRNA-seq techniques has now enabled individual groups to measure dozens or even hundreds of samples. Ongoing consortium efforts are also underway to generate extensive atlases of single-cell datasets covering diverse biological contexts with thousands of samples^8,9^. Joint consideration of such panels poses additional technical and conceptual analysis challenges. The panels can be extensive in terms of the number of samples and diverse in terms of their composition, with some of the datasets lacking any shared cell subpopulations. Panels will often encompass distinct categories of samples, such as treatment / control sets, samples with disease and normal pathology, or different tissues. In these cases, the desired behavior of the joint analysis method will depend on the question being posed by the investigator. For example, in examining a panel of normal and metastasis-infiltrated tissue samples^10,11^, it may be desirable for a method to identify a subset of exhausted CD8^+^ T lymphocytes as a distinct cluster for a detailed follow up. However, the investigator may instead wish to contrast the state of such cells with their closest counterpart in the normal tissue, in which case it would be desirable for a method to identify a cluster of cytotoxic CD8^+^ T cells spanning both normal and metastasis samples. We therefore set out to develop a flexible approach for analyzing and navigating such sample collections.

We reasoned that a unified graph representation could be used to capture likely relationships between cells in different samples, and that statistical analysis of such a joint graph can identify subpopulations forming across different samples at different scales (Figure 1a). To construct the joint graph, Conos performs pairwise alignments of samples within the panel to identify plausible inter-sample cell-cell correspondence (inter-sample edges). Such mappings are expected to be error-prone. For instance, if the two samples being compared do not share any subpopulations, any established edges between such samples will be erroneous. Across many such pair-wise comparisons, however, the recurrent subpopulations of cells will tend to map to each other, forming cliques within the joint graph that can be identified as cell communities over the background noise of spurious edges (Figure 1a). To improve statistical power, Conos also adds low-weight edges connecting neighboring cells within individual samples, which serve as a weak prior for preserving local neighborhoods of cells in each sample. A variety of techniques can be used for establishing a plausible mapping between a pair of samples. Inter-sample edges can be established simple strategies such as nearest-neighbor or mutual nearest neighbor (mNN) mapping between samples^7^. The similarity between cells in different samples can be assessed using reduced space that would minimize the impact of sample-specific variation. For instance, Buttler *et al.* showed that calculating neighbors in the space of canonical variables results improves mapping between analogous samples^6^. We focused on joint matrix factorization approaches that identify projections capturing common variation across two or more samples, including common principal component analysis^12^ (CPCA), joint non-negative matrix factorization (JNMF), and higher order generalized singular value decomposition^13^ (HO GSVD).

**Figure 1.**
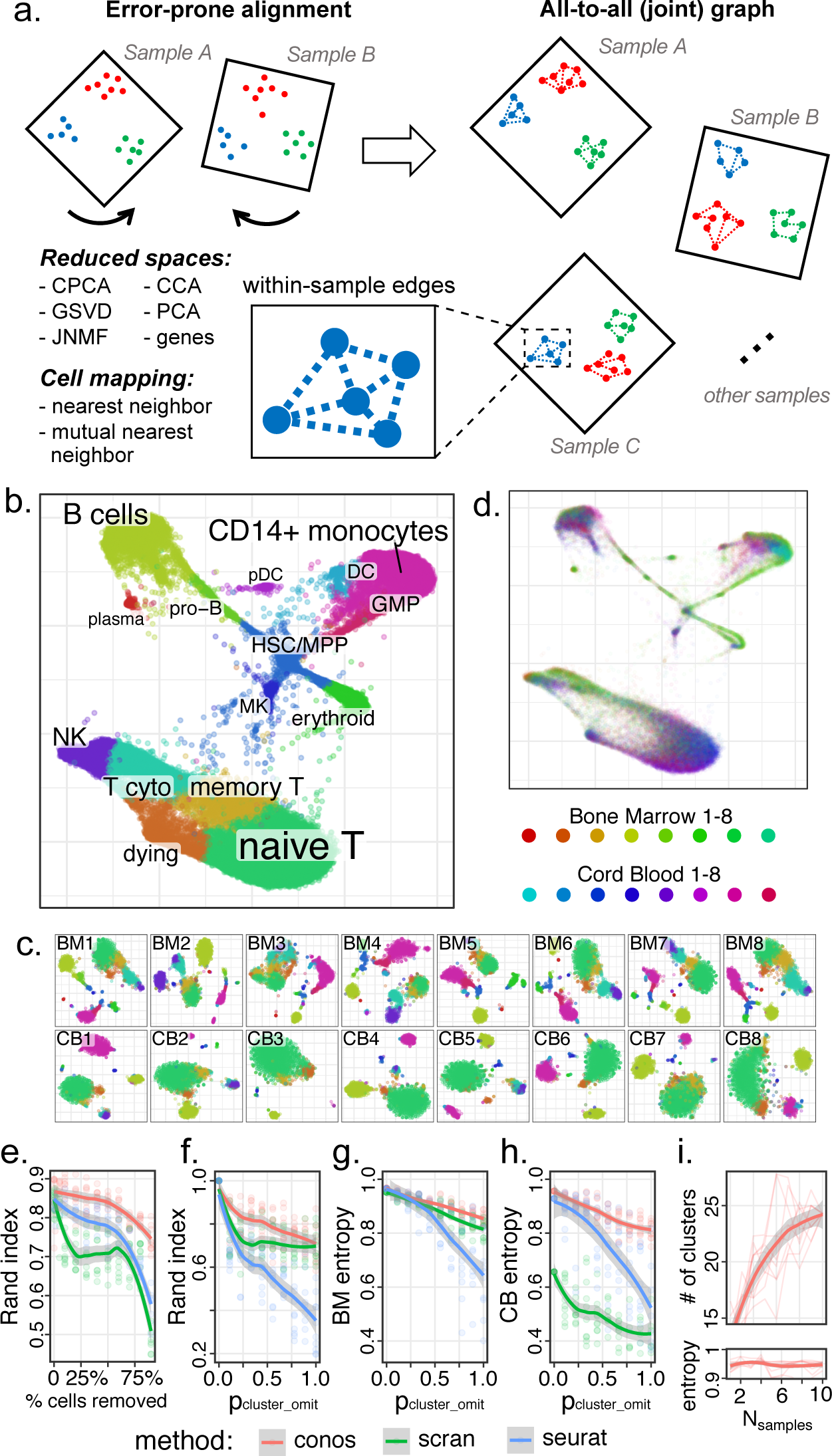
Joint graph is an effective strategy for assembling diverse scRNA-seq dataset collections. **a.** Conos establishes initial mapping between datasets based on common transcriptional variation, as determined by CPCA, JNMF, or alternative factorizations. In a heterogeneous collection, such mapping is expected to be error prone. By comparing many pairs of datasets, Conos builds a joint graph whose primary edges connect potential neighbors across datasets (inter-sample edges), with additional low-weight edges connecting neighbors within individual datasets. Subpopulations of cells recurrent within the dataset collection form clique-like communities of inter-sample edges within the joint graph. **b.** Joint graph combining eight human bone marrow and eight cord blood datasets is visualized using largeVis embedding, labeling major cell types (see Supp. Figure 1 for details and an alternative embedding). **c.** Mixing of cells from different datasets in the joint graph. The graph retains systematic distinctions between tissues and individual samples that show significant deviation in their composition that can be analyzed further. **d.** Communities detected on the joint graph provide unified clustering throughout the collection, as illustrated on individual dataset t-SNE embeddings. **e.** Adjusted Rand index (y-axis) is shown as a function of the fraction of cells omitted from the datasets (x-axis) relative to the full dataset for different joint clustering approaches. Conos shows improved stability of subpopulation detection even for small numbers of cells. **f.** Stability of the subpopulation detection is shown for increasing amount of heterogeneity between datasets. Adjusted Rand index is shown for increasing probability of random subpopulation omission from individual datasets (x-axis, see Methods). **g,h.** Mixing of different bone marrow (h) and cord blood (i) datasets within the identified subpopulations is quantified using normalized average cluster entropy (see Methods). **i.** The power to detect cell subpopulations increases with the size of the collection. The number of stable clusters (y axis, see Methods) detected in a collection of human bone marrow samples increases as more samples are added to the collection (x-axis), while maintaining high level of sample mixing (high average cluster entropy) within each cluster.

We first examined the performance of Conos approach using a combination of CPCA and mNN mapping on a collection combining eighteen scRNA-seq samples of human bone marrow and cord blood from the Human Cell Atlas^9^. Projection of the resulting joint graph separated all major subpopulations (Figure 1b, Supp. Figure 1), with the detected joint cell clusters connecting the corresponding subpopulations across the entire collection (Figure 1c). While the individual samples were well-mixed within the joint graph and the resulting clusters, the bias in the composition of the two tissues was also apparent, with stem cell and progenitor populations as well as cytotoxic T cells being more abundant in the bone marrow, while naïve T cells or megakaryocytes being more prevalent in cord blood (Figure 1d). To quantify robustness and sensitivity of different approaches we perturbed the full data using strategies that decreased signal or increased heterogeneity between samples. Examining the ability to recover the original subpopulations identified by each method under decreasing numbers of cells (Figure 1e) or under decreasing magnitude of subpopulation-specific expression signatures (Supp. Figure 1e), we find that CPCA in combination with mutual nearest neighbor mapping shows most robust performance, significantly outperforming earlier methods. Conos performance remained robust even for the simple pairwise alignment strategies (nearest neighbor mapping based on simple gene correlation), highlighting the power of the joint graph approach to community detection. We have applied Conos to re-analyze a number of recently published complex datasets ^11,14-16^, in all cases joining corresponding annotated subpopulations across different samples and tissues (Supp. Figures 2-7).

Modern experimental designs are likely to combine distinct classes of samples within the panels, such as sets of disease samples and healthy controls, or multiple tissues from different individuals. We simulated increasing heterogeneity of a panel by omitting increasing number of random clusters from different samples, and evaluating the method’s ability to recover originally detected subpopulations (Figure 1f). Conos showed higher robustness compared to other methods. Furthermore, Conos was able to sustain uniform mixing (*i.e*. high entropy) of cells from different samples among the identified joint clusters even for very high degree of sample heterogeneity, where some of the samples shared few or no common subpopulations (Figure 1g,h). Community detection on a joint graph shows high sensitivity, for instance enabling Conos to detect subpopulations that may be represented by only a single cell in a given sample by linking them to corresponding sets of cells in other samples (Supp. Figure 1f). Detection of such rare or subtle subpopulations can be greatly improved when one or more samples in the panel happens to contain a more sizeable representation of this subpopulation, facilitating formation of the corresponding community structure in the joint graph. More importantly, the sensitivity of the proposed approach increases as more samples are added to the panel (Figure 1i), suggesting that increasing collections of scRNA-seq samples will reveal more subtle or rare cell subpopulations.

Consideration of diverse sample panels requires one to re-examine the aims of alignment and identification of joint clusters within the panel. While for homogeneous panels, the analysis simply aims to identify a broad set of clusters that appear in nearly every sample, this is not the case for panels including diverse classes of samples. For example, when analyzing a panel containing both tumor and adjacent normal samples^16^, it would not be desirable for a method to lump tumor cells with any normal tissue subpopulations, even though the cluster of tumor cells would be restricted to a subset of samples (*e.g.* Supp. Figure 5). In a more nuanced scenario, the clustering may separate tumor-associated CD4^+^ T cells from their counter-parts in the healthy tissues^14^, picking up persistent biological difference in their state (Figure 2a-d). However, if the investigator aims to compare the transcriptional shifts associated with the state of the CD4^+^ T cells in the tumor environment, it would be more suitable to identify a unified CD4^+^ T cell cluster spanning both tumor and adjacent normal samples (Figure 2e). Graph communities can be viewed as a hierarchical clustering of cells, and in that way the difference between separating or joining tissue-specific subgroups of CD4^+^ T cells is equivalent to cutting the hierarchy at different levels (Figure 2d, Supp. Figure 5).

**Figure 2.**
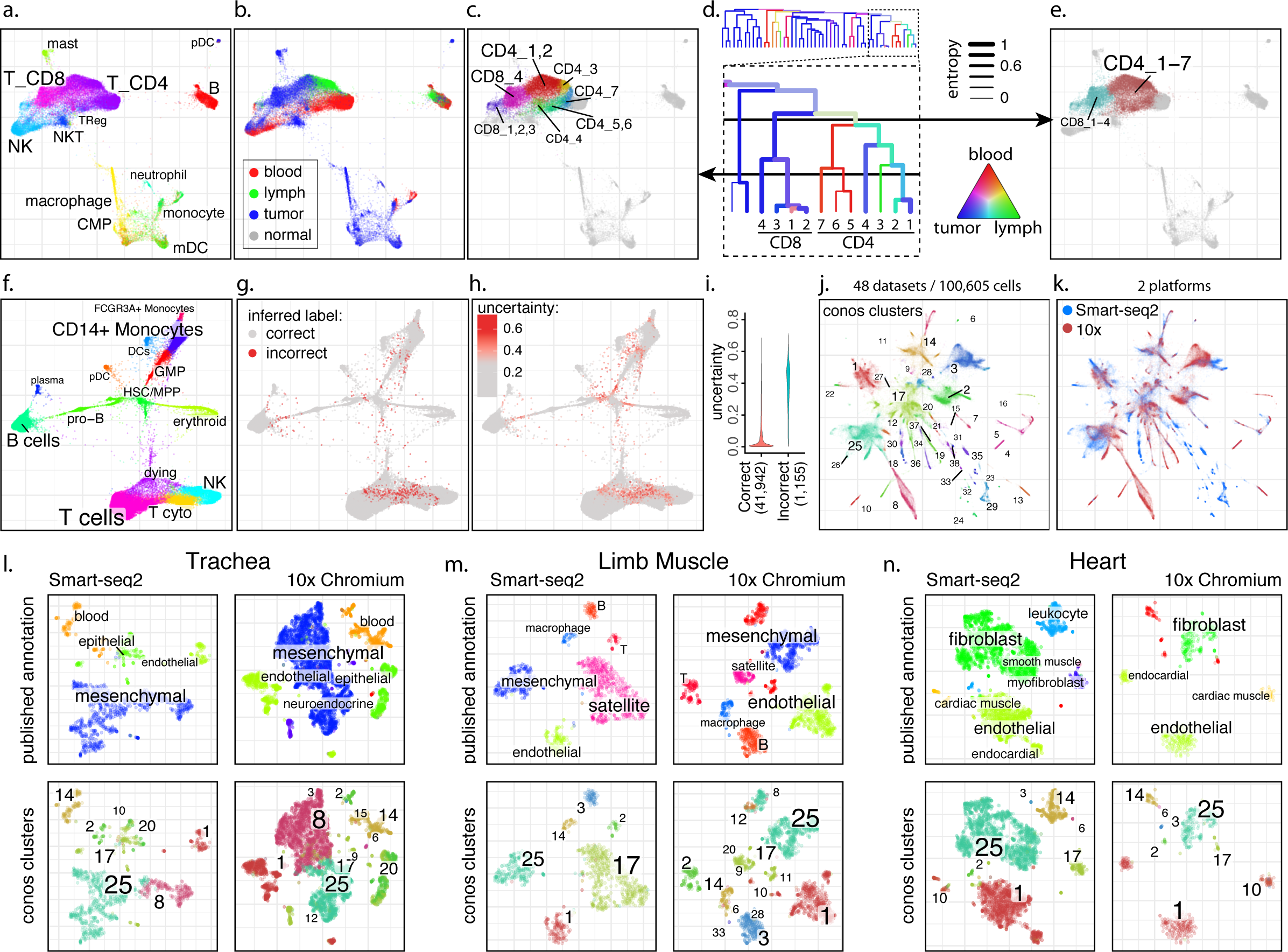
Examples of dataset analyses using joint graphs. **a-e.** Trade-off between cluster resolution and sample breadth. An embedding of a joint graph combining immune microenvironment cells from eight breast cancer patients^14^ is shown (a). The distribution of source tissues within the joint graph (b) highlights systematic expression differences in T-cell and some monocyte populations between tissues. A fragment of the subpopulation hierarchy is shown for T cells subsets (d), with color of the branches showing tissue composition, and width showing normalized sample entropy (higher entropy corresponds to more samples being in the branch). Depending on the level, a cut of the cluster hierarchy can yield more focused but tissue-specific clusters (c) or less granular clusters that bridge across more tissues and samples (e). **f-i.** Propagation of cell annotation labels. Joint embedding of bone marrow samples from eight patients is shown (f), annotating major subpopulations. The annotations were erased from all but one sample, and propagated back to the entire dataset. The labels were inferred correctly for 97% of the cells, with incorrect inferences (g) occurring near the cluster borders or at poorly-covered cells. These coincided with cells for which Conos reported high uncertainty of the labeling (h,i). **j-k.** Conos integration of the Tabula Muris dataset^8^. Joint graph of the 48 separate datasets covering different mouse tissues is shown (100,605 cells), with colors and numbers marking top-level joint clusters (j). The collection contains samples measured using different scRNA-seq platforms, which are well mixed within the shared subpopulations (k). Comparison of the joint clusters with the published annotation is shown for three tissues (l-n) that were measured with both platforms. Joint clustering shows consistency between tissues and platforms, with some clusters giving higher resolution (*e.g.* separation of blood or mesenchymal populations in trachea samples), and others joining related cell types across tissues (*e.g.* fibroblast and part of the mesenchymal population are joined under cluster 4 in l-n).

In general, cutting lower in the tree will yield higher resolution of subpopulations – such as ability to distinguish more subtle subsets of monocytes. On the other hand, lower cuts are also likely to decrease cluster breadth – the average fraction of samples in the panel in which a cluster appears. As the balance between the desired resolution and breadth will depend on the question being posed by the investigator, Conos implements an interactive tool to explore the hierarchical community structures, such the one shown in Figure 2d. For a given panel, the web application allows investigators to choose progressively deeper cuts, while monitoring breadth and distribution of the resulting clusters, as well as the extent to which different subsets of samples in the panel are separated by the composition of the resulting clusters (Supp. Figure 8).

Diffusion of information on the joint graph provides a natural way of mapping properties across samples. For instance, one can propagate discrete values, such as cell annotation labels to datasets that have not yet been annotated. To demonstrate this, we asked Conos to propagate cell cluster labels from one human bone marrow dataset to the remaining seven unlabeled datasets (Figure 2f). By propagating existing labels through the graph using diffusion process, and keeping track of the uncertainty, Conos is able to label cells with 97% accuracy (Figure 2g). The mislabeled cells were mostly found along border separating adjacent clusters, and were reported to have high uncertainty of the labels (Figure 2h,i). Graph diffusion can also be used to propagate contiguous values. Existing batch correction methods have generally aimed to coerce multiple datasets into a common gene expression space, typically by adjusting gene expression values of all cells to eliminate all perceived sample-specific (batch) effects. Conos estimates such adjusted expression values by diffusing the cell-specific expression estimates through the graph (Supp. Figure 9). Alternatively, if reduced-dimensional representation of the common expression space is desired, Conos can achieve this quickly by starting diffusion with cell projections on the Laplacian eigenvectors of the joint graph. We contend, however, that in most settings, where it cannot be assumed that the differences between samples are caused predominantly by measurement noise, the utility of such “batch-adjusted” expression values will be mostly limited to visualization. Once the appropriate clustering of cells is established on the panel, we expect follow up analyses to focus on the expression variation among samples. This includes subpopulation-specific analysis of expression differences between groups of samples, or variation within such groups.

Overall, our results demonstrate that integration of extensive single-cell collections into a unified graph representation can provide effective means for integrative analysis of such data, including subpopulation identification, differential expression or annotation. Compared to existing methods, Conos implementation shows improved stability and sensitivity, particularly notable on heterogeneous sample panels, such as multi-tissue / multi-patient clinical study designs. This robustness allows Conos to wire together diverse collections of samples, such as organism-scale atlases where most of the samples will have few or no cell types in common. For example, we reanalyzed a recently published mouse atlas^8^, combining 48 datasets covering different mouse tissues (Figure 2j). Conos was effective at identifying common cell populations across samples measuring diverse tissues, as well as overcoming the differences of the two different scRNA-seq platforms utilized (Figure 2j-n, Supp. Figures 10-12). The approach is relatively fast, particularly when using a simpler PCA space (*e.g.* integration and clustering of the 100,605 cell mouse atlas took less than two minutes on a dual-processor machine), though analysis of much larger collections will likely require to limit the number of pairwise sample comparisons. Further gains can be made with improved community detection algorithms, or by using iterative statistical analysis of the graph structure to identify more informative inter-sample edges. We hope that the presented approach and its further developments will enable other research groups to effectively explore and apply complex single-cell RNA-seq experimental designs.

## Methods

### Overview of the approach

Conos processing can be divided into several key phases. During the **phase I**, each individual dataset in the sample panel is filtered and normalized using standard packages for single-dataset processing: pagoda2 or Seurat. Specifically, Conos relies on these methods to perform cell filtering, library size normalization, identification of overdispersed genes, and in case of pagoda2 - variance normalization. During **phase II**, Conos performs pairwise comparisons of the datasets in the panel to establish initial, error-prone, mapping between cells of different datasets. These inter-sample edges are then combined with lower-weight intra-sample edges during **phase III** – joint graph construction. The joint graph is then used for downstream analysis, including community detection, label propagation, *etc*.

### Pairwise dataset alignments (phase II)

Initial inter-sample edges between a given pair of datasets *i* and *j* was established based on a choice of *1*) rotation space, and *2*) neighbor mapping strategy (nearest neighbor or mutual nearest neighbor). For each dataset, a set of overdispersed (hypervariable) genes (*g^i^*, *g^j^*) was determined using pagoda2 (by default n.odgenes=2000 top overdispersed genes were used), and a union of overdispersed genes from both datasets was taken, limiting the genes to those for which data was available in both datasets: *g* = [*g^i^* ∪ *g^j^*] ∩ *G^i^* ∩ *G^j^*, where *G^i^* and *G^j^* are the total gene sets for the two datasets. Subsequent analysis was carried out on matrices *M^i^* and *M^j^*, with columns of the matrix correspond to genes *g*, and rows correspond to cells of each dataset. The entries of each matrix were taken to be variance-normalized expression magnitudes (as determined by pagoda2).

Reduced projection matrices *R^i^* and *R^j^* were obtained according to the space used: For CPCA and JNMF, these corresponded to projections onto the corresponding common/joint components (30 components, by default). For PCA space, PCs (50 by default) were determined independently for *M^i^* and *M^j^*, with *R^i^* and *R^j^* then determined by projecting cells of each dataset to a joint (*i.e.* 100-dimensional) space of both PCs. For gene space, *R^i^* and *R^j^* were taken to be matrices *M^i^* and *M^j^* themselves.

Cell-cell similarity between cells *k* and *l* were determined as *w_kl_* = *r_kl_*, where *r_kl_* is the Pearson linear correlation between the *k*-th row of *M^i^* and *l*-th row of *M^j^*. An alternative L2 distance was implemented as 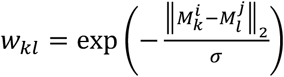, where the default scaling constant *σ* = 10^5^.

### Joint graph (phase III)

For a given sample collection, the nodes of the joint graph *G* corresponded to all of the cells included in the collection, connected by a combination of inter- and intra-sample edges. The inter-sample edges were determined as mutual nearest neighbors (mNN, default) or plain nearest neighbors (NN), with the weight *w_kl_*. Neighborhood size k=15 was used by default. Intra-sample edges for each dataset *i* connected each cell to *k_self_* (default *k_self_* = 5) cells within the dataset *i* using weights *w_kl_* = *c_self_* · *r_jk_* within the reduced space *R^i^* as determined by the projection of cells on the top PCs of *M^i^*. Constant *c_self_* = 0.1 was used to reduce the contribution of the intra-sample edges relative to the inter-sample edges. For visualization purposes, joint graphs were laid out in two dimensions using *largeVis* algorithm varying parameter *alpha* between 0.5 and 2.5, depending on the complexity of the dataset.

### Joint clustering

Joint clusters were determined as communities of the joint graph *G*, using standard community detection methods. By default, *walktrap.communities* algorithm implemented by the *igraph* package was used, with step=20. Louvain clustering implemented by *igraph::multilevel.communities* method provided much faster performance, but lacked hierarchical output.

### Label and value propagation

Propagation of both labels and expression magnitudes was treated as a general problem of information propagation between graph vertices. Graph vertices can have multiple labels, either continuous or discrete. Such labels can be affected by biases or different kinds of noise. Assuming that adjacent vertices on the graph have similar labels, we can reduce this noise using iterative diffusion process on the joint graph. For continuous labels the diffusion process was implemented as follows:

1. At the beginning of an iteration, each vertex has a label *L_i_*; An edge between vertices *i* and *j* has a weight *c_i,j_*.
2. During the iteration, for each label we update its value with 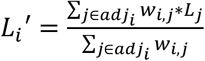, where *w_i,j_* = *exp*(−*a*(*c_i,j_* + *b*), *a* and *b* are hyper-parameters of the algorithm (default values *a* = 10, b=0.5 were used). The set *adj_i_* includes all vertices, adjacent to *i*, including vertex *i*(*w_i,i_* = 1).
3. Infinity norm of difference between the two labelings (*max_i_*|*L_i_*′ − *L_i_*|) is used as the termination condition.

Considering each gene as a continuous label, Conos uses this process to correct gene expression matrices to bring all the datasets into a “common” expression space. We note that a single iteration of the diffusion process with parameter *a=0* is equivalent summing of expression over adjacent cells, which is a common approach for noise correction in scRNA-seq data.

For discrete labels, the implementation tracked label uncertainty, with the diffusion process being used to estimate posterior probability of each label for each vertex. This was performed by running diffusions on the probability distribution of the labels:

1. Posterior distribution of possible labels was kept for each vertex. The starting state for the labeled vertices was set so that the probability of the true label is set to 1, with the probability of other values set to 0. For the unlabeled vertices, all of the values were initially set to 0.
2. On these distributions we simultaneously run diffusion process for each component of the distribution (*i.e.* for each label). After each diffusion step, the values of the posterior distributions were re-normalized so that the sum of the values was equal to 1.

The diffusion of discrete values was used for the cell annotation propagation results (Figure 2f-h). In the figures, the uncertainty of the labeling was evaluated 1 − *max_i_*(*p_i_*).

### Benchmark design

Quantitative performance of different methods shown in Figure 1 and Supp. Figure 1 relied on the same general design, where each method *m* was ran on a full dataset to obtain clustering *C^m^*. The full dataset was then gradually perturbed to pose a more challenging problem, and the ability of different methods to recover their corresponding original clustering *C^m^* was measured. Such was aimed to place different methods on equal footing, and make use of realistic data (as opposed to synthetic). Details of different benchmarks are given below:

- Cell subsampling benchmark (Figure 1e). HCA BM+CB 3k dataset containing a total of 16 samples was used (see below for dataset details). A percentage of cells *p_removed_* ∈ [0, 80%] (x-axis) was randomly sampled and removed from each dataset in the collection. For each value of *p_removed_*, a total of 10 replicates of dataset perturbation were generated. To assess performance, adjusted Rand index (y axis) was calculated relative to the first replicate with *p_removed_* = 0. A smoothed mean for each method and the corresponding 95% confidence band is shown on the Figure 1e were calculated using *igraph::geom_smooth()* method. Note that all of the examined methods show certain level of instability to negligible perturbations of the dataset (such as shuffling of the cell order in the matrix, or removal of a single cell). As datasets sampled with *p_removed_* = 0 shuffled the order of the cells, the adjusted Rand index value at *p_removed_* = 0 is below 1.
- Cell mixing benchmark (Supp. Figure 1e). HCA BM+CB 3k dataset was used. For each dataset *i*, background expression vector *b^i^* was determined for each gene *g* as 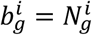, where 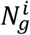 is the total number of molecules of gene *g* detected in the dataset *i*. A perturbed dataset with a mixing proportion *p_mix_* ∈ [0,1] was generated for each cell by iterating through each molecule of the cell, keeping the original molecule with a probability 1 − *p_mix_*, or alternatively (with probability *p_mix_*) replacing it with a molecule randomly sampled from the background profile *b^i^*. This way datasets generated with *p_mix_* = 0 are equivalent to the original data, whereas *p_mix_* = 1 yields datasets where each cell is a random sampling of the background, and any cell subpopulations would be impossible to discern.
- Cluster dropping benchmark (Figure 1f-h). To simulate increasing compositional variability between samples in the collection, cells belonging to a cluster *c* ∈ *C^m^* were omitted with a probability *p_omit_*. The sampling procedure was carried out independently for each dataset, so that different subsets of clusters were dropped from different datasets (increasing compositional variability). To guarantee a minimal dataset size, a total of 5 clusters were sampled this way in each dataset. Under such procedure, *p_omit_* = 0 maintains the full original dataset, whereas *p_omit_* = 1 maximizes inter-sample compositional differences. The degree to which cells from different samples were mixed within the resulting clusters was quantified using normalized entropy, weighted by the cluster size:

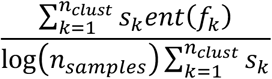

where *f_k_* is a vector giving the number of cells from each sample in a cluster *k, ent()* is the entropy (calculated using *entropy* package in R), *s_k_* is the total number of cells in a cluster *k, n_clust_* is the number of clusters detected by the method on a current realization of the dataset, and *n_samples_* is the total number of samples in the panel. Given that some systematic composition differences between bone marrow (BM) and cord blood (CB) are expected, normalized entropy was assessed separately for BM and CB cells (Figure 1g and 1h, respectively).
- Number of stable clusters (Figure 1i). To assess how the number of stable clusters changes with the increasing size of the sample panel, we assembled a larger panel of samples covering the same tissue (HCA+10x BM dataset). Ten randomized “series” were constructed, with each series starting with two randomly chosen datasets, and then adding one dataset per step up to a maximum of 10 available datasets (sampling without replacement was used to construct the series). As community detection algorithms rely on heuristics such as maximization of modularity, we evaluated the number of stably detectable clusters as a number of independent subtrees in the hierarchy returned by the *walktrap.community* algorithm. A stable subtree was determined as a subtree containing at least 30 cells that can be detected under a 10% cell subsampling perturbation (see below) with sensitivity and specificity above 0.8. To evaluate these stability properties, for each run additional 10 subsampling runs were made omitting 10% of the cells of the sample and rerunning *walktrap.community* to generate perturbed trees.
- Sensitivity to individual cells (Supp. Figure 1f). To evaluate how well different methods are able to pick up rare cells in the dataset, we simulated rare cell occurrences by randomly choosing a single sample in the panel, then choosing a random joint cluster *c* ∈ *C^m^* that occurs within that dataset and leaving only one randomly selected cell from that cluster *c* within that sample. HCA BM+CB 1k panel was used, containing 16 samples. A total of 16 × 5 × 5 = 400 perturbed panels were generated, sampling five different clusters *c* from each of the 16 datasets, with five different random choices of the remaining cell being made. To evaluate the performance, the remaining cell from the cluster *c* was scored as correctly classified if it was assigned to a cluster to which other cells of a cluster *c* were most commonly assigned.

### Implementation of other methods

Conos performance was compared with the two previously published methods, configured in the following way:

- Seurat package was installed from CRAN. The pre-processing and dataset alignment was ran as recommended in the tutorial: http://satijalab.org/seurat/immune_alignment.html
- The mNN approach by Haghverdi *et al*. was ran by installing *scran* package from CRAN. Hypervariable genes for each dataset were determined according to the tutorial. To enable execution on large datasets within the available memory constraints, the number of hypervariable genes was limited to the top 2000 genes (same number as used for Conos), based on the sum rank of genes across dataset-specific hypervariable gene lists. To determine joint clustering, an approach analogous to Conos was used: k-nearest neighbor graph (k=30) was constructed based on the 30 top PCs of the adjusted expression values, and *igraph::walktrap.community* method was used to identify cell clusters.

To enable large-scale benchmarking, the number of common space components estimated by Conos and the two methods above was limited to 20.

### Data sources and dataset-specific analysis details

1. Human Cell Atlas (HCA) bone marrow and cord blood was downloaded from the HCA portal (https://preview.data.humancellatlas.org/). To reduce calculation times in benchmark evaluations, we took a random subset of the cells from lane1 of each dataset. 3000 cells per sample were used by default (HCA BM+CB 3k datasets). A smaller, 1000 cell dataset (HCA BM+CB 1k) was used for the more extensive sensitivity analysis (Supp. Figure 1f).
2. For Figure 1i, we combined HCA BM samples with two samples (“Frozen BMMCs Healthy Donors 1 and 2”) downloaded from 10x Genomics (https://www.10xgenomics.com/resources/datasets/).
3. Azizi *et al.* data on breast cancer was downloaded from GEO as a count matrix, together with the provided annotations. In showing the plots (Figure 2, Supp. Figure 4) the annotations were simplified to collapse patient-specific populations and omit smaller subpopulation distinctions. To demonstrate applicability to different levels of data fragmentation, the dataset was re-analyzed by combining either 8 patients, 15 patient+tissue combinations, or 53 patient+tissue+replicate combinations.
4. Lambretchs *et al.* molecular count data and annotations on the lung cancer were downloaded from ArrayExpress.
5. Guo *et al.* molecular count data and annotations non-small-cell lung cancer were downloaded from GEO (GSE99254).
6. Puram *et al.* molecular count data and annotations on head-and-neck cancer were downloaded from GEO (GSE103322).
7. Human cortex count matrix for Hoghe *et al.* bioRxiv 2018 was downloaded from downloaded from http://celltypes.brain-map.org/rnaseq. Lake *et al.* count matrix was downloaded from GEO.
8. Tabula Muris mouse data was downloaded from https://tabula-muris.ds.czbiohub.org/. Only cells with at least 1000 molecules were analyzed. A total of 48 datasets were combined.

### Availability

Conos is implemented as an R package with C++ optimizations, and is available on GitHub (https://github.com/hms-dbmi/conos) under the GPL-3 open source license.

